# Rapid design and prototyping of biocatalytic virus-like particle nanoreactors

**DOI:** 10.1101/2022.02.10.479872

**Authors:** Lygie Esquirol, Donna McNeale, Trevor Douglas, Claudia E Vickers, Frank Sainsbury

## Abstract

Protein cages are attractive as molecular scaffolds for the fundamental study of enzymes and metabolons, and for the creation of biocatalytic nanoreactors for *in vitro* and *in vivo* use. Virus-like particles (VLPs) such as those derived from the P22 bacteriophage capsid protein make versatile self-assembling protein cages and can be used to encapsulate a broad range of protein cargos. *In vivo* encapsulation of enzymes within VLPs requires fusion to the coat protein or a scaffold protein. However, the expression level, stability and activity of cargo proteins can vary upon fusion. Moreover, it has been shown that molecular crowding of enzymes inside virus-like particles can affect their catalytic properties. Consequently, testing of numerous parameters is required for production of the most efficient nanoreactor for a given cargo enzyme. Here we present a set of acceptor vectors that provide a quick and efficient way to build, test and optimise cargo loading inside P22 virus-like particles. We prototyped the system using yellow fluorescent protein then applied it to mevalonate kinases, a key enzyme class in the industrially important terpene (isoprenoid) synthesis pathway. Different mevalonate kinases required considerably different approaches to deliver maximal encapsulation as well as optimal kinetic parameters, demonstrating the value of being able to rapidly access a variety of encapsulation strategies. The vector system described here provides an approach to optimise cargo enzyme behaviour in bespoke P22 nanoreactors. This will facilitate industrial applications as well as basic research on nanoreactor-cargo behaviour.

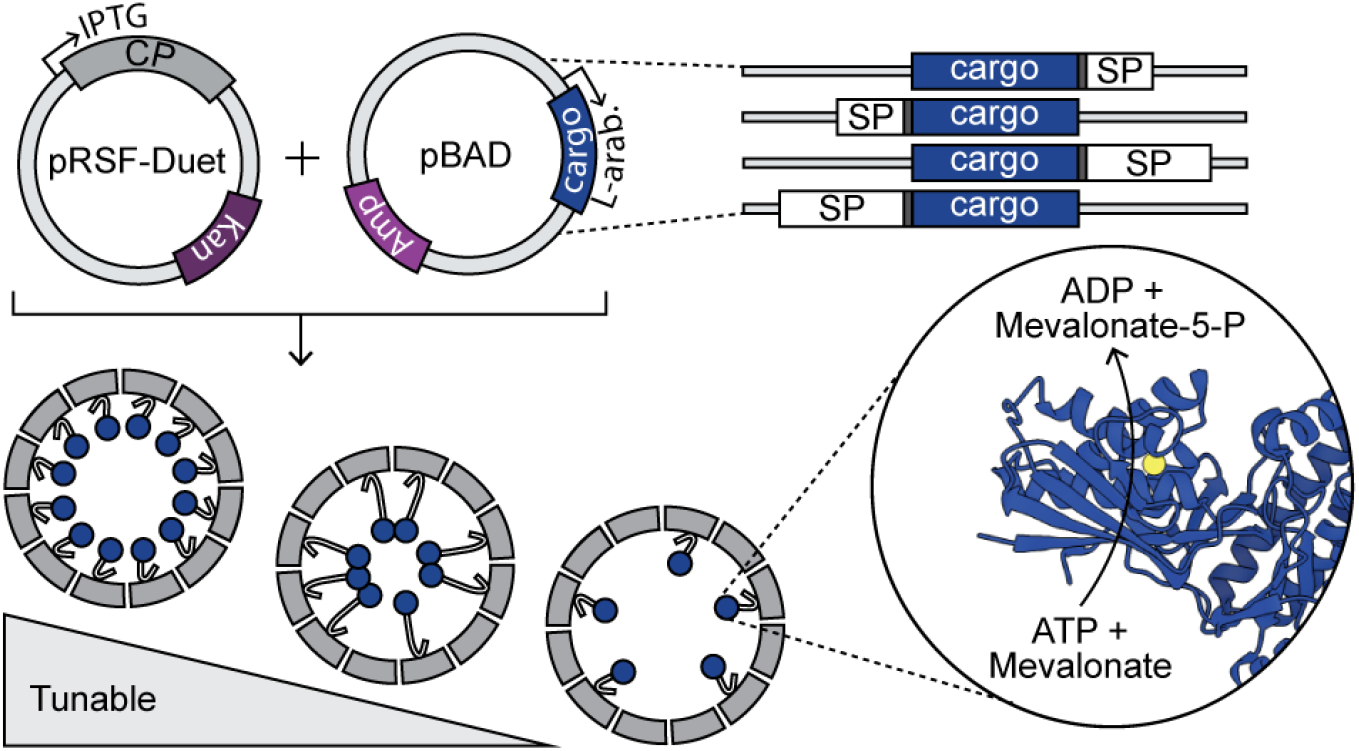

## INTRODUCTION

Compartmentalization is a hallmark of cellular life. In Eukaryotes, membrane-bound organelles allow for segregation of enzymes that would otherwise be competing for similar substrates, creation of microenvironments with different pH and salt concentration, and the segregation of toxic compounds within the cells. In prokaryotes, compartments are micro-structures made of proteins instead of membranes. Examples include the encapsulins, which have been observed across a range of microorganisms [1] and are thought to play a role in stress resistance [2]; and carboxysomes in cyanobacteria, which trap and concentrate CO_2_ around the RuBisCO enzyme [3] resulting in one of the most efficient RuBisCO enzymes amongst photosynthetic organisms [4]. Viruses are also nano-scale compartments that operate *in vivo*, albeit typically to the disadvantage of host cells. Virus-like particles (VLPs) derived from the structural proteins of viruses are examples of engineered nano-scale compartments that can be used as bespoke nano-reactors. Encapsulation of enzymes by heterologous protein cages such as VLPs can improve resistance to proteases, chaotropes, and thermal denaturation [5-8]. These protective properties, combined with the flexibility to encapsulate multiple enzymes [9-11], have increased interest in VLP engineering as nanocompartments for applied *in vitro* [12,13] and *in vivo* [7,8,14,15] biocatalysis in manufacturing and health.

VLPs of the *Salmonella typhimurium* Bacteriophage P22 have emerged as a robust and efficient system for assembling tightly packed biocatalytic nanoreactors in *E. coli* [16,17]. The P22 capsid is composed of 420 copies of the coat protein (CP) that assemble with the help of 100-300 scaffold proteins (SPs) into a 55 nm icosahedral nanoparticle [18,19]. By fusing a protein cargo to the SP, or simply to the portion of SP that interacts with the inner surface of the capsid [20], the fusion partner is encapsulated during capsid assembly [16]. P22 VLPs offer great potential for co-localisation of small metabolic pathways [9] and biocatalytic nanoreactors assembled from P22 VLPs have been engineered to enable the production of molecular hydrogen [12]; and developed as hierarchical, responsive biomaterials with enhanced catalytic properties [13,21].

As a tool for fundamental research, the P22 platform also provides a convenient compartment to study the impact of confinement and crowding on enzyme kinetics. Within cells, protein concentration is in the range of 100-300 mg/ml [22] – a concentration that is normally not possible to achieve *in vitro*. Molecular crowding is known to impact numerous metabolic processes, from enzyme catalytic rates to protein folding [23] and investigating these processes has largely relied on the use of synthetic crowding agents. While not reproducing the complex environment of a cell, VLPs provide a tool enabling very high ‘cell-like’ protein concentrations of encapsulated proteins [24,25], allowing the study of enzyme behaviour in crowded environments. Using VLPs to concentrate and confine enzymes has led to intriguing effects on protein stability [26] and catalytic properties [11]. A generalizable yet not fully understood impact is one of increased thermal stability. This has been observed for numerous enzymes encapsulated within VLPs and suggests that these crowded environments inhibit protein unfolding [6,8].

Understanding these emergent properties to the point where they can be intentionally and specifically implemented requires control over enzyme loading densities. While decreased catalytic efficiency of encapsulated enzymes has been often observed [16,27], the negative impact of crowding on the *k*_*cat*_ of alcohol dehydrogenase (AdhD) was only shown by encapsulating different densities of AdhD inside P22 VLPs using *in vitro* assembly [10]. In this example, VLPs were assembled with control over the stoichiometry of separately isolated coat protein and cargo protein. However, programmable loading during *in vitro* assembly requires high purity of components [28,29] and is, therefore, not a cost effective strategy at scale. *In vitro* assembly approaches can help identify optimal cargo densities for biocatalytic nanoreactors, but to investigate and understand the impacts of confinement on enzymatic activities more broadly, and in the context of *in vivo* biocatalysis, *in vivo* assembly of biocatalytic nanoreactors with tuneable loading densities is required.

Here we present a method to rapidly test different parameters that influence expression, encapsulation, and enzyme behaviour in the P22 VLP platform. The system allows tuneable control of multiple parameters and is suitable for high throughput analysis of combinatorial cargo protein designs *in vivo*. The orientation of enzyme fusions are known to impact activity [30]; the relative timing of the expression of enzyme cargo and coat protein can be an important factor in enzyme maturation before encapsulation in VLPs [12]; and the loading density is expected to be influenced by both the timing of expression and the level of cargo enzyme induction. The system presented here allows for facile testing of these parameters to optimise the influence of loading density on enzyme activities within P22-based biocatalytic nanoreactors. The mevalonate pathway supplies precursors for the biosynthesis of over 70,000 terpenes from all domains of life, including numerous industrially important compounds [31]. Phosphorylation is central to the pathway and the first phosphorylative step is catalysed by mevalonate kinase. Using three orthologues of this key class of enzyme we show that the optimal configuration for loading density and catalytic efficiency is enzyme-dependent, highlighting the need for a streamlined platform to test multiple conditions.

## RESULTS AND DISCUSSION

### System Design

To enable efficient discovery of optimal conditions for *in vitro* biocatalysis or as novel *in vivo* metabolic engineering tools [14,32,33], the system should be amenable to high-throughput DNA assembly. We therefore built a set of “acceptor” vectors that can be used for quick Golden Gate-based construction and subsequent testing of differentially loaded P22 VLPs. For any given gene, eight constructs can be derived from a single donor clone, each with a combination of the following parameters: SP length, orientation of the SP fusion, and vector backbone allowing for simultaneous or independent induction of coat and cargo (Figure 1). Using the L-arabinose inducible promoter also allows tuning of cargo expression levels [34]. The acceptor vectors have been designed to allow quicker cloning, relying on an enhancement of the Golden Gate methodology [35] that utilises the lethal *ccdB* gene to select against non-recombined acceptor vectors [36,37].

**Figure 1.**
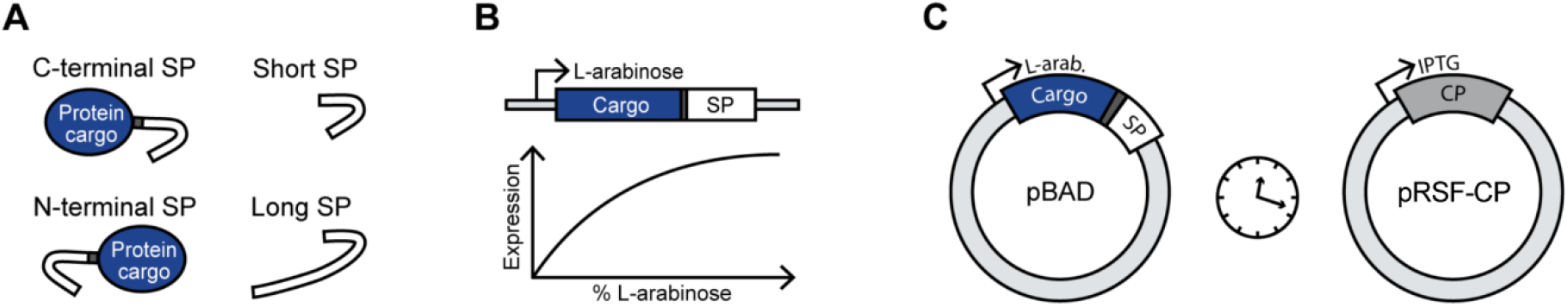
Options for each of the parameters that influence cargo loading by P22 VLPs. (A) The orientation and length of the SP fusion to the cargo protein, (B) tuning expression levels using the L-arabinose promoter pBAD, and (C) control over the timing of the expression with two different promoters allowing differential expression of the SP-cargo fusion and CP.

Both the acceptor and the donor vectors are flanked with compatible sites produced by the Type IIs restriction enzyme *BbsI*. Simultaneously applying the restriction enzyme and T4 DNA ligase in one pot reactions with a donor plasmid carrying the gene of interest and individual acceptor plasmids results in swapping of excised BbsI fragments between the two (Figure 2). Acceptor vectors contain either kanamycin (pRSFDuet-1 backbone) or ampicillin (pBAD backbone) resistance cassettes. When using the system, the acceptor vector should contain a different resistance cassette from the donor vector (which may have, for example, a chloramphenicol resistance cassette). Combining the Golden Gate approach and the *ccdB* lethal selection system resulted in extremely high cloning efficiency. Out of a total of 154 colonies sequenced from 61 constructs made to date with this cloning system (includes constructs outside of this study), 149 were found to be correct, making the cloning efficiency of the recombination reactions 97%.

**Figure 2.**
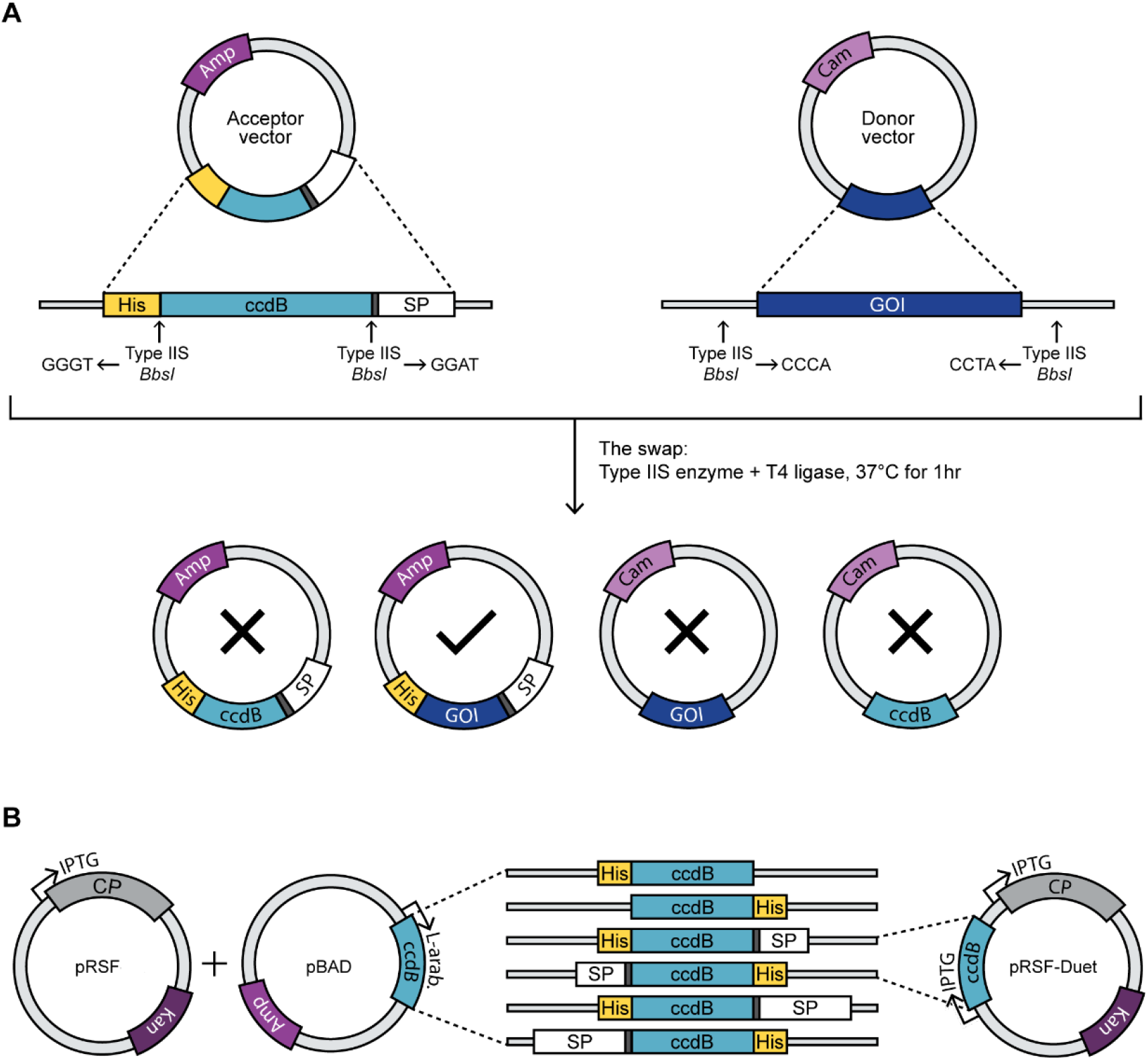
Principle of the cloning strategy and available acceptor vectors. A) Generalised acceptor vector containing a ccdB lethal cassette (left) and a donor vector containing a gene of interest (right), each flanked by complementary type IIS cloning sites. The matching cohesive ends result in four possible ligation outcomes vectors, but only one is viable on appropriate antibiotic selection as indicated by a tick. B) Different ccdB cassettes contained in our six pBAD and two pRSFDuet-1 acceptor vectors. SP: scaffold protein, His: 6X His-tag, GOI: gene of Interest; SP: Scaffold protein, CP: coat protein, linker (grey).

A one-hour isothermal recombination reaction is performed for each acceptor vector from a single donor plasmid (Figure 2A). The system allows for quick cloning of one gene of interest into 8 different acceptor vectors. Six of these comprise the pBAD backbone for L-arabinose tuneable expression of the cargo protein as His-tagged SP fusions or a His-tagged protein with no SP fusion for purification independent of the VLP system (Figure 2B). For translational fusions of the cargo protein to truncated SP there is a choice of either the N-terminus or C-terminus and, for each of these, SP variants of two different lengths are available comprising residues 141-303 (long SP) or residues 238-303 (short SP). The pBAD-based expression vectors are compatible with the pRSF plasmid that supplies the P22 coat protein (CP) under an IPTG-inducible promoter, which allows for independent induction of the CP and SP fusions. Co-expression of any of SP-fused cargo protein and the CP results in encapsulation of the cargo protein in a P22 VLP. The final two acceptor vectors use the bicistronic expression cassette from pRSFDuet-1 for the simultaneous expression of a short SP-fusion at the N or C terminus along with the CP. These last two acceptor vectors were not used in this study but are available for use along with the other vectors and all plasmid sequences are provided in the Supplementary Data.

### Differential VLP cargo loading can be achieved *in vivo* by varying construct design

We first parameterised the vector series using yellow fluorescent protein (YFP). Acceptor vectors containing short SP, located either at the N- or C-terminus of the cargo YFP in the pBAD backbone, were used. When co-expressed with the pRSF CP plasmid, the SP fusion drives encapsulation of the cargo protein during assembly of the VLP *in vivo*. Factors influencing VLP loading in this experiment are the orientation of SP fusion and the strength of L-arabinose induction of the SP fusion. Comparing high (0.2%) and low (0.004%) L-arabinose resulted in a significant variation in loading density in purified VLPs with the higher L-arabinose concentration resulting in higher loading for both SP orientations (Figure 3A). In addition, the orientation of the fusion influenced loading density at both induction levels, with the N-terminal SP resulting in higher loading, and with this effect considerably greater for the high induction level. We observed a 6-fold variation between lowest and highest loading, from 45 to 275 cargo proteins per VLP (Figure 3B), resulting in internal SP-YFP concentrations ranging from 1.4 mM to 8.7 mM (equivalent to approximately 50 to 300 mg/mL). Despite the large apparent difference in the number of scaffolding proteins per VLP, this does not have a noticeable effect on VLP morphology (Figure 3C), consistent with previous observations [9,12,25,29]. The structural consistency is a hallmark of the system and ensures that the organisation of the protein shell and, therefore, the pores for substrate, co-factor, and product diffusion are likely to also remain constant. This is particularly useful for investigating the possible role of molecular crowding and confinement on enzyme function, as the available volume of each nanoreactor remains the same, while the density of the cargo varies [10].

**Figure 3.**
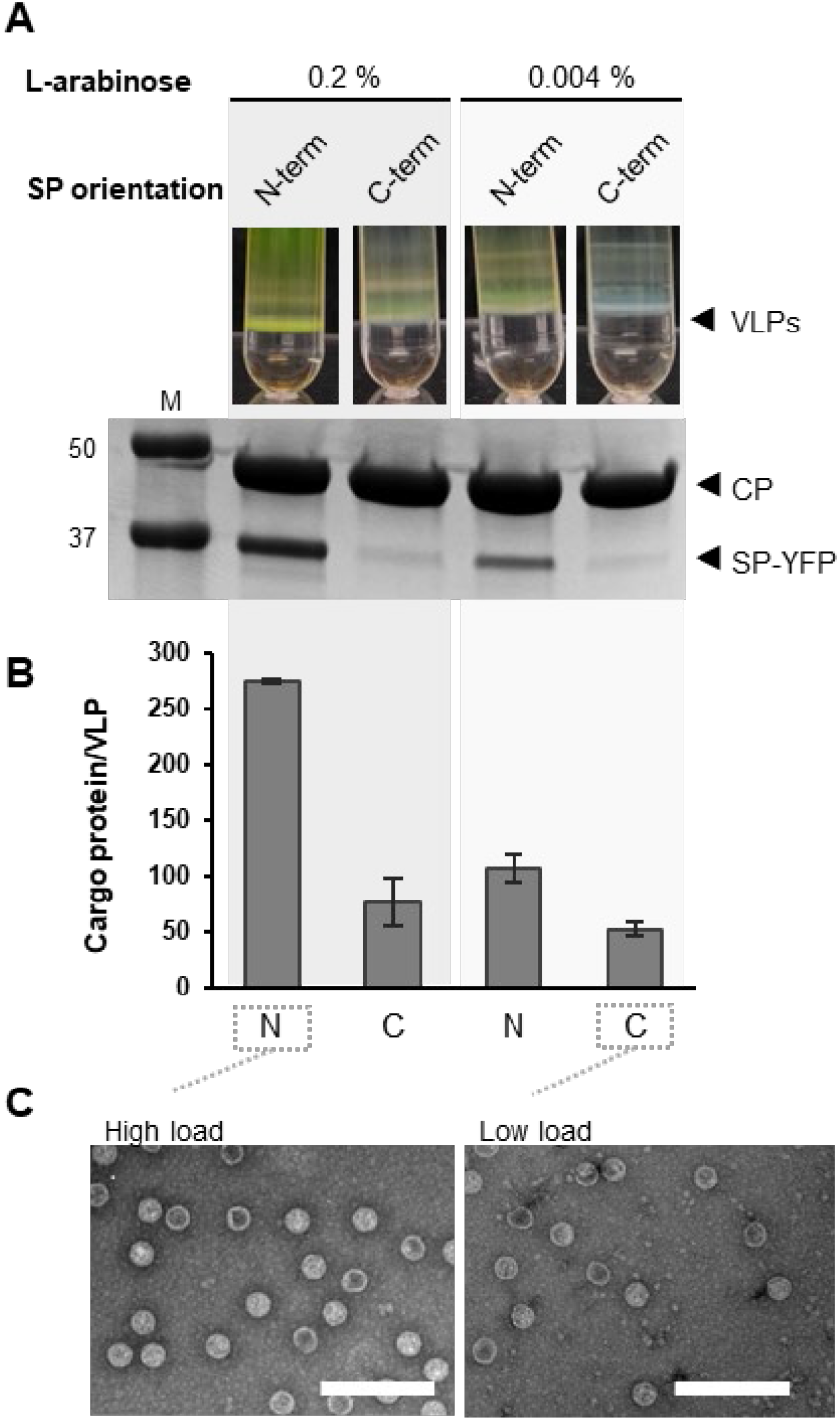
Impact of SP orientation and induction strength on YFP loading in P22 VLPs. A) Ultracentrifugation with iodixanol gradient results in a band containing P22 VLPs (black arrow) and SDS-PAGE of the VLP band obtained under different expression conditions (fusion either in N-terminus or C-terminus, and either high (0.2%) or low (0.004%) arabinose induction. B) Densitometric analysis of YFP loading inside P22 VLPs. C) Representative transmission electron micrographs of P22 particles showing VLP morphology under high loading or low loading conditions.

### VLP loading is idiosyncratic and depends on both the encapsulation strategy and the cargo

We next applied our system to the cloning and encapsulation of the three mevalonate kinases (MKs). MK is a key enzyme of the mevalonate isoprenoid pathway [38]; it plays an important role in both native pathway flux and in metabolic engineering [31,39]. The mevalonate pathway is highly conserved and exists across all domains of life [40]. It is responsible for production of over 70,000 different metabolites from cholesterol to hormones, many of which have useful industrial applications [31,41,42]. Deficiency in MK has been linked to autoinflammatory disorders in humans such as periodic fever syndrome, hyperimmunoglobulinemia D, and mevalonic aciduria [43]. Moreover, some MKs are regulated by feedback inhibition of prenylphosphate compounds synthesised downstream of the mevalonate pathway [44]. MK is therefore an interesting target in need of greater understanding, both in human health and for metabolic engineering applications [45].

We chose three MK orthologues to investigate the optimisation of VLP confinement: one from the psychrophilic archaeon *Methanococcoides burtonii* (MKbur) [46]; one from the tardigrade *Ramazzottius varieornatus* (MKvar) [47] (both previously uncharacterised); and the structurally and functionally characterised MK from the archaeon *Methanosarcina mazei* (MKmaz) [48]. These enzymes add the first of two phosphates to the mevalonate pathway biochemistry: they use ATP to phosphorylate mevalonate, forming mevalonate-5-phosphate (Scheme 1). Phosphates are key to isoprenoid biochemistry, since the energy inherent in phosphate bonds is used to drive many of the downstream reactions used for structural diversification and, ultimately, biological functionality. All three MKs were assembled into constructs for fusion with both long and short versions of SP at either the N-terminus or the C-terminus. Each set of strains co-expressing an SP-MK fusion and the CP construct was induced at high (0.2 %) and low (0.004%) L-arabinose, resulting in eight MK-VLP test systems for each enzyme.

**Scheme 1.**
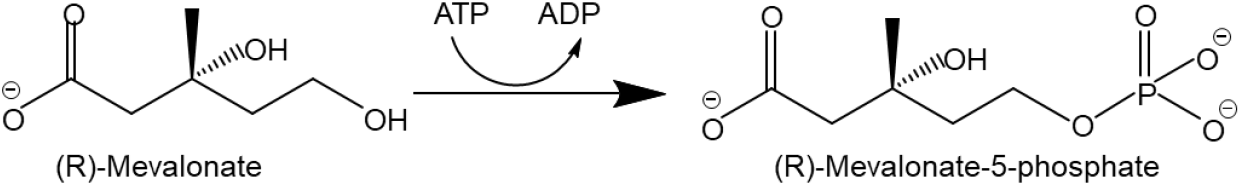
Reaction catalysed by mevalonate kinases.

As observed with the YFP encapsulation (Figure 3), higher induction strength generally resulted in higher cargo loading for MKs (Supplementary Figure 1). On the other hand, SP orientation and SP length had orthologue-dependent impacts on MK loading (Supplementary Figure 1).To determine whether there were statistically significant effects on loading density for each of the expression and SP fusion parameters, we analysed each parameter across all cargo proteins (Figure 4). The loading level was most significantly impacted by the concentration of arabinose used for expression (P-value 0.00002) (Supplementary Table 2). However, when looking at individual cargo proteins, impact of the concentration of arabinose on loading density was only statistically significant for YFP. The next most significant influence on loading density was the identity of the cargo protein (P-value 0.0003) (Supplementary Table 2). Accordingly, the impact of both orientation and length of the SP fusion was variable, orthologue dependent, and in some cases interacting. Orientation had a non-significant though noticeable impact on the loading overall (p-value 0.07; Supplementary Table 2) with significantly different loading only observed for YFP (Figure 4). The N-terminal fusions tended to lead to higher loading than the C-terminal fusions for MKmaz, MKbur and YFP (Figure 4), however for MKvar the opposite can be observed with the C-terminal fusion leading to higher loading (Figure 4). Alignment of the MKs show considerable differences in the N-terminal part of the sequence for MKvar (Supplementary Figure 3), which could result in differences in the way various components of the system interact. The length of the SP fusion had a statistically significant impact on loading overall (p-value 0.00007, Supplementary Table 2). This parameter was not tested for YFP as that cargo was only encapsulated using the short SP. However, for both MKmaz and MKbur the impact of the scaffold length was statistically significant with opposite effects of the different lengths for each orthologue; long SP for MKmaz and short SP for MKbur (Figure 4).

**Figure 4.**
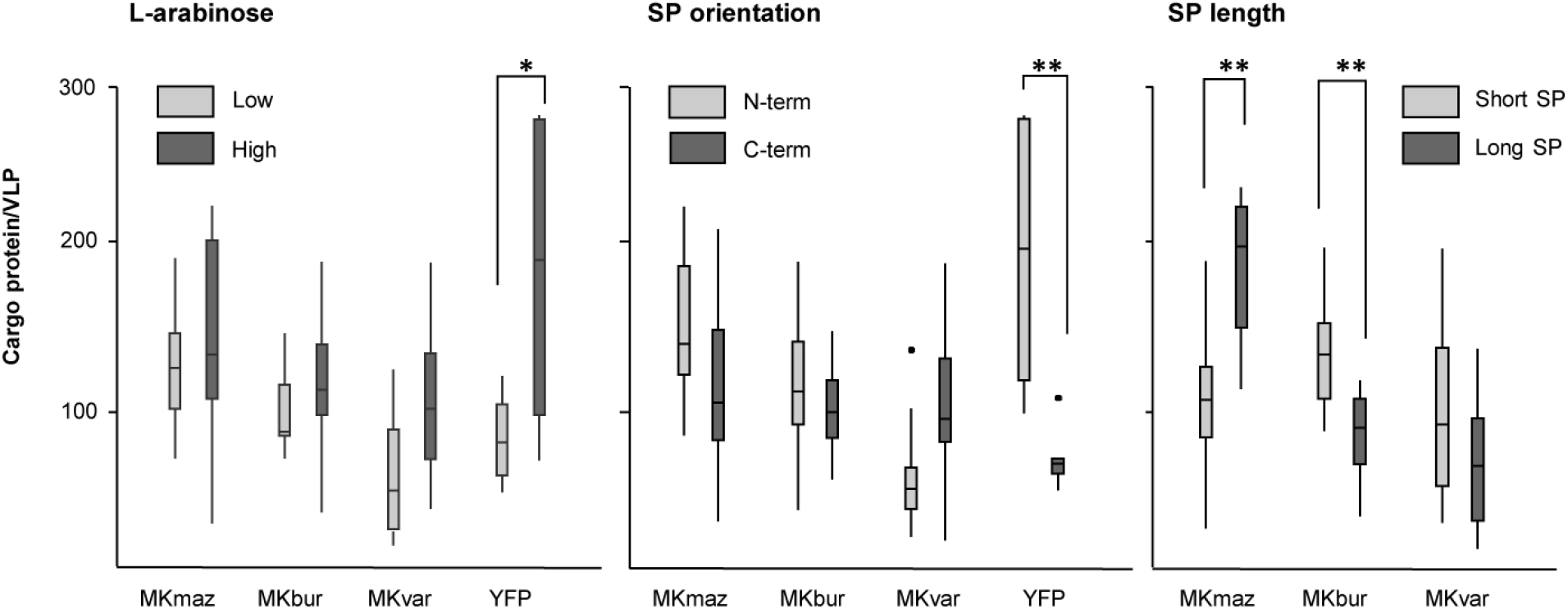
The impact different expression and SP fusion parameters have on loading for individual cargo proteins. Individual boxplots for VLP loading depending on the amount of arabinose used for cargo expression (high = 0.2%, low = 0.004%); the SP orientation (N-term or C-term); and the fusion length (long or short scaffold protein). Statistically significant differences for each protein cargo are shown where ‘*’ and ‘**’ represent p-values of 0.05 and 0.01, respectively.

In summary, differential loading of VLPs was obtained *in vivo* using combinations of different fusion and expression parameters. Overall, induction strength, which is expected to result in differential expression level, offers a general mechanism of extrinsic control over loading. On the other hand, the impact of SP orientation and length, which are intrinsic protein engineering approaches, are highly cargo dependent. This mirrors the finding of cargo-specific impacts on expression and encapsulation into Qβ *in vivo* [5]. That work, as well as the data presented here, illustrates the challenges of predicting the impact of encapsulation in complex systems such as *in vivo* assembling VLPs. Here, we show that ostensibly similar proteins (Supplementary Figure 2) have remarkably different loading levels in response to different sequence contexts. Sequence-specific phenomena, including the fidelity of folding, stability of the SP fusion, variability at termini, and ability of the SP fusion to direct VLP assembly are all likely contributing factors. Differential timing of SP-cargo and CP induction was not tested in the present study and is also likely to be influenced by these protein specific factors. Together these results highlight the need for screening design parameters for each cargo protein candidate and the utility of our system for the straightforward and rapid assembly of a series of constructs to optimise *in vivo* encapsulation in terms of loading density.

### The impact of VLP loading strategy and density on kinetic properties is enzyme-specific

Both theoretical models and empirical studies clearly demonstrate that the highest loading does not lead to the highest enzyme efficiency in VLPs [24,49]. It is therefore critical to be able to control loading density if VLPs are to be used as a tool for creating effective biocatalytic nanoreactors. Kinetic properties were determined for MK-VLPs of each MK orthologue to analyse effects of the encapsulation strategy on encapsulated enzyme kinetics compared to their free and SP-fused counterparts. Enzyme efficiency was chosen as an illustration of MK-VLP performance as it allows performance comparisons between different enzymes. Enzyme efficiency is calculated by dividing the enzyme’s turnover *k*_cat_ (number of molecules of substrate converted into product per second) by the enzyme’s *k*_M_ (its ability to bind to its substrate). Since the enzymes were assayed using a coupled assay, taking indirect measurements of MK activity, we report the apparent kinetic parameters. All Michaelis-Menten curves (Supplementary Figures 3-5) and kinetic parameters (Supplementary Tables 2-4) for enzymes and various VLPs can be found in the supplementary information.

To examine the effect of N- or C-terminal SP fusions on enzyme behaviour in the absence of encapsulation, the free and SP-tagged MKs were expressed and purified in the absence of the CP construct. The efficiency of the free MKs was quite similar, with an apparent *k*_*cat*_ /apparent *k*_M_ of 82.1, 87.5 and 92.3 for MKmaz, MKbur, and MKvar, respectively (Figure 5). Our kinetic data for free MKmaz (Supplementary Table 2) is in agreement with previous publications, which report an apparent *k*_M_ (app *k*_M_) of 0.12 mM and an apparent *k*_cat_ (app *k*_cat_) of 10.1 sec^-1^ [48,50,51]. As reported for other MKs originating from the archaea [48,50], no substrate or product-mediated inhibition was observed for MKmaz or MKbur (Supplementary Figures 3 and 4). However, inhibition of MK plays a central role in the regulation of the terpenoid pathway in most organisms [44,52,53] and MKvar was found to be inhibited above 3 mM of substrate, with a K_i_ of 2.7 mM (Supplementary Figure 5), consistent with substrate inhibition previously reported for an MK from *Staphylococcus* [53]. For each MK orthologue, SP fusion reduced the efficiency of MKs relative to their unfused free counterparts with the effect being variable, depending on the orthologue as well as the orientation and length of SP (Figure 5). SP fusion decreased efficiency by 40-70 % for MKmaz, 25-75 % for MKbur, and 80-90 % for MKvar. For MKmaz and MKbur, the N-terminal fusion performed better than the C-terminal fusion; and for MKvar the opposite was the case. For MKbur, the short SP fusion performed better; for the other two enzymes, length of the SP did not have a strong effect. Polypeptide fusions are commonly found to negatively impact enzyme kinetics [54], so these results are not entirely unexpected, although the effect on MKvar is particularly detrimental.

**Figure 5.**
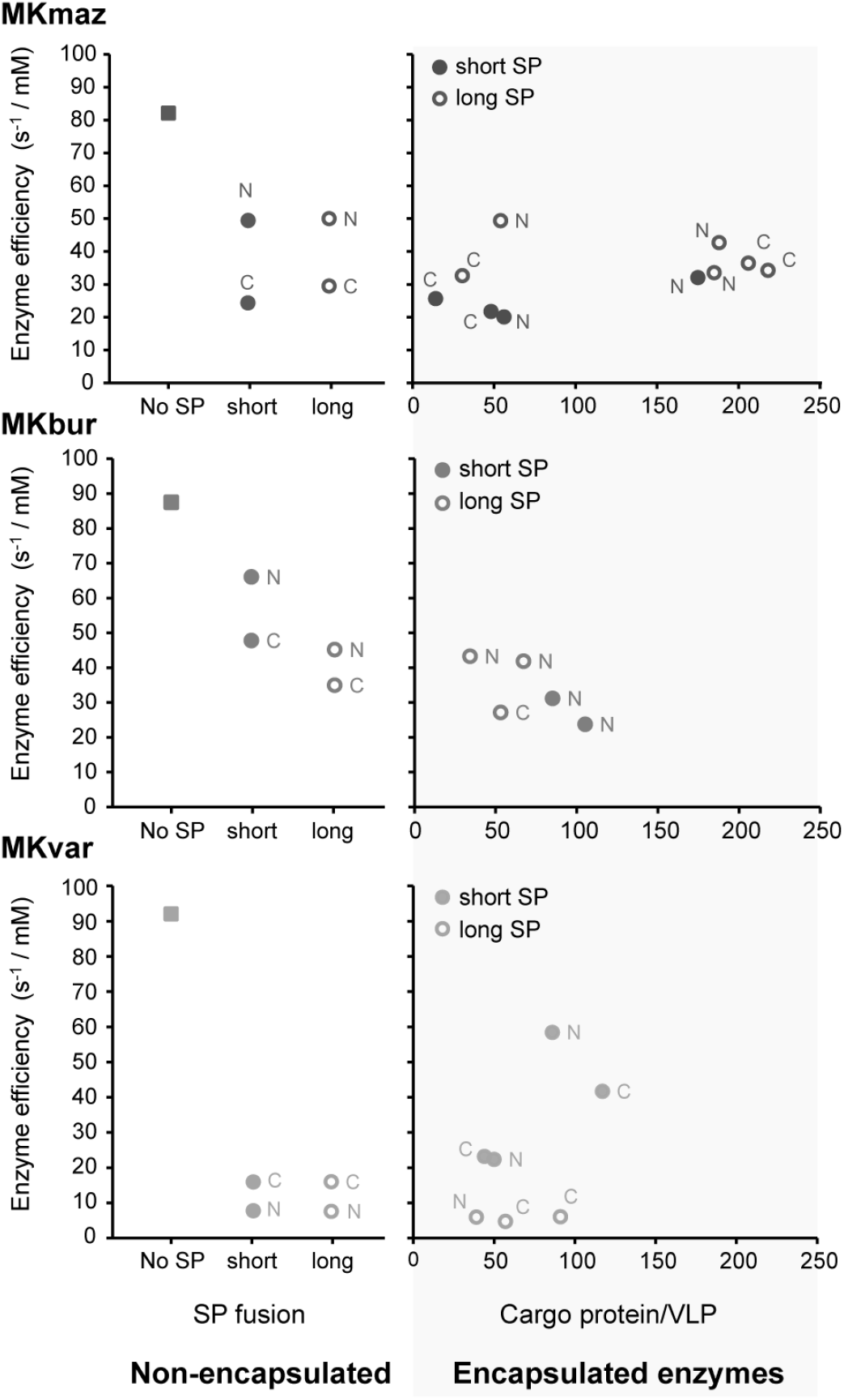
Enzyme efficiencies calculated for MKs, SP-MK fusions, MK-VLPs. MKmaz = dark grey, MKbur = mid-grey, MKvar = light grey. Low-loading MKmaz replicates highlighted with a dashed outline.

To investigate the impact of VLP encapsulation on the efficiency of MK-VLP nanoreactors we characterised 10 MKmaz VLPs (with loading ranging from 14 to 218 MK per VLP), 7 MKvar VLPs (with loading from 39 to 117 copies), and 5 MKbur VLPs (with loading from 34 to 105 copies). For MKmaz we were able to analyse both high- and low-loading MKmaz VLPs, but there was no apparent effect of enzyme loading on the efficiency of this orthologue (Figure 5). In contrast to the free SP-tagged MKmaz, there was no difference between C- and N-terminal fusions for the encapsulated enzyme and in general encapsulation did not have much of an effect on efficiency when compared to the free enzymes. For MKbur VLPs, encapsulation of short SP fusions had a negative impact on efficiency whereas the long SP fusions, which were less efficient when unencapsulated, were not affected by loading into VLPs (Figure 5). For MKvar, encapsulation of short SP fusions significantly improved efficiency compared to encapsulation of long SP fusions (p-value <0.05) and compared to the free SP-tagged enzyme, recovering some of the efficiency lost in the SP fusion. For the slightly higher loaded examples, there was an efficiency improvement of 3-fold to 4-fold over the free SP fusion (Figure 5). There was no apparent effect of SP orientation. In summary, for these MK orthologues we do not see a clear relationship between enzyme efficiency and any of the tested variables besides the noted impacts of encapsulation.

In this study we compared VLP-based nanoreactors to both free enzymes and to the enzymes fused to the obligatory anchor molecule that directs their encapsulation, the SP. Fusion to an SP reduced catalytic efficiency for all MK orthologues tested, with the effect being dramatic in the case of MKvar (Figure 5). Both the long and the short SP are largely disordered [55] and carry high positive charges (pI of short SP = 10.9; pI of long SP = 9.9). It is therefore possible that the fusion directly interferes with the active site in addition to a more general effect on fusion partner structure [54]. Orthologue-specific impacts of SP fusions on MKs is found in the differential impacts of fusion orientation on loading densities (Figure 4) and catalytic efficiency (Figure 5). However, recovery of efficiency upon encapsulation was observed for some MKvar VLPs. Interaction of the C-terminal helix turn helix of SP with CP [56] could reorient SP away from the cargo enzyme, mitigating the negative impact of SP fusion on *k*_*cat*_. This phenomenon was observed for the short SP fusions of MKvar, which displayed an average 8.3-fold higher app *k*_*cat*_ (Supplementary Table 4), significantly different to long SP fusions across all loading densities (p-value <0.001). Interestingly, encapsulation also slightly alleviated the inhibition of MKvar, raising the *k*_i_ (Supplementary Table 4). Overall, we observe a highly idiosyncratic response to SP fusion and encapsulation among MK orthologues. This points to a need to empirically determine the optimal encapsulation strategy and loading density for biocatalytic nanoreactor assembly. While the efficiency of the enzymes decreased upon SP fusion and, for the most part, encapsulation, for certain applications other benefits of encapsulation might counteract this loss. For example, the increased higher thermostability [6] and stabilisation of unstable enzymes [7,26] that has been observed upon encapsulation.

## CONCLUSION

We have produced a set of vectors for fast construction, testing and optimisation of P22-based biocatalytic nanoreactors via enzyme fusion to a domain of the capsid scaffolding protein. A simple one-step recombination reaction will produce a set of vectors combining different parameters that control the expression and encapsulation of cargo proteins. Our results show that effect of SP length and orientation as well as induction level have strong effects on VLP loading density and that the effect of individual parameters are unpredictable and cargo specific. Moreover, we show that these encapsulation parameters affect the activity of encapsulated enzymes in an enzyme-dependent manner with some influence of loading density. The enzymes tested here were three orthologues of a key enzyme in the mevalonate pathway. The disparate effects of encapsulation parameters on their activity underscore the need to empirically optimise biocatalytic nanoreactor construction.

Our data highlight the difficulty of achieving predictable behaviours in proteins, which is an aim of synthetic biology and which has been achieved to a much greater extent for DNA-based components such as promoters and terminators [57,58]. They also reflect the utility of having access to combinatorial construction tools [59], which provide the opportunity to increase the examinable solution space and optimise for the desired outcome. The use of recombinant proteins shells that emulate natural compartments is growing in interest for both biomedical [60,61] and manufacturing applications [17,62]. The system presented here provides an accessible methodology to optimise the performance of biocatalytic nanoreactors in a comprehensive yet streamlined manner for the popular and well-characterised P22 VLP platform.

## MATERIALS AND METHODS

### Molecular cloning

#### Acceptor vectors

Cassettes used for building acceptor vectors were designed and ordered as dsDNA fragments (gBlocks) from Integrated DNA Technologies (IDT). The different elements used to build the GoldenGate cassettes are built around a central *ccdB* gene flanked by outward facing *Bbs*I sites resulting in unique 5’ (GGGT) and 3’ (GGAT) overhangs. Outside of the *Bbs*I sites is a sequence encoding a 6X His-tag sequence at either the 5’ or 3’ end with, optionally, a sequence encoding for a short glycine/serine linker leading to either a short (amino acids 248-303) or a long (amino acids 141-303) C-terminal portion of the P22 scaffold protein at the opposite end. The full set of arrangements results in the six different acceptor vector cassettes (Figure 1) which were inserted into the pBAD vector backbone via Gibson assembly cloning downstream of the ARA promoter. Similarly, GoldenGate cassettes were inserted by Gibson assembly downstream of the first T7 promoter of a pRSF-Duet vector already containing the P22 CP following the second T7 promoter [27]. In this case, the lacI gene was simultaneously replaced with a gBlock encoding a GoldenGate domesticated version to remove the two unwanted *Bbs*I sites. The pRSF-Duet vectors were not used in this study, however, the nucleotide sequences for all acceptor vectors as well as the complementary pRSF-P22CP construct are provided in Supplementary Data and the plasmids will be made available from Addgene. The acceptor vectors were cloned and maintained using *ccd*B Survival 2 T1^R^ cells (Thermofisher.com).

#### Other vectors used and genes of interest

A PCR product of the pRSF backbone and the P22 CP expression cassette was amplified from the aforementioned pRSF-Duet source vector [27] and re-ligated by homology directed repair to create the monocistronic pRSF-P22CP plasmid (Supplementary Data). Donor vectors containing our genes of interest were ordered from Twist biosciences (USA) as clonal genes in the pTwist high copy plasmid with chloramphenicol resistance. The sequences were designed to flank the gene of interest by two inward facing *Bbs*I sites leaving the nucleotides GGGT in 5’ and GGAT in 3’, matching the acceptor vector’s overhangs respectively (Figure 2). Sequences used in this publication encode: EYFP, Mevalonate kinase (MVK1) from *Methanosarcina mazei* (WP_011033702.1, Q8PW39) [48], an uncharacterised Mevalonate kinase (MVK3) from the psychrophilic archaeon *Methanococcoides burtonii* (WP_011500381.1, Q12TI0) [46], an uncharacterised Mevalonate kinase (MVK5) from *Ramazzottius varieornatus* (BDGG01000012.1, GAV05667.1) [47].

#### Recombination into the acceptor vectors

The GoldenGate-mediated exchange of the *ccdB* cassette with genes of interest was performed by mixing 50 ng of acceptor vector (pBAD carrying ampicillin resistance or pRSF carrying kanamycin resistance) and 50 ng of donor plasmid with 1 µL *Bbs*I (NewEngland biolabs, USA) and 0.25 µL T4 ligase (NewEngland biolabs, USA) in a 20 µL total volume. The mixture is incubated at 37 °C for one hour, before transformation into Dh5-alpha competent cells. The counter-selection will lead to a very low number of incorrect clones as any *ccdB* containing vectors will be lethal for standard cloning strains such as Dh5-alpha, thereby eliminating uncut acceptor vectors, and selection for antibiotic resistance carried by the acceptor vectors will eliminate chloramphenicol resistant donor vectors.

### Protein expression and purification

#### Free proteins

For the His-tag-mediated purification of free and fused enzymes recombined pBAD acceptor vectors were used to transform Escherichia coli BL21 (λDE3) cells. Bacteria were grown on Luria-Bertani (LB) medium containing 100 μg/mL ampicillin at 37°C, shaking at 180 rpm until OD_600_ reached 0.6. Induction was triggered by addition of 0.2% L-arabinose and the temperature was lowered to 28°C. Cells were incubated over night before their harvest by centrifugation at 5,000 x g for 15 minutes. Cells were resuspended in Buffer A (100nM NaCl and 5 mM imidazole in 50 mM Tris at pH 8.0) and lysed by passage through a homogenizer Emulsiflex-C5 (AVESTIN, Canada) five times at 20,000 PSI. Lysis was followed by centrifugation at 18,000 x g for 30 minutes to pellet the cellular debris, and the supernatant was filtered through 0.22 µm syringe filter (Millipore, USA). The soluble fraction was then loaded onto a His-Trap column (GE Healthcare Life Sciences) and eluted with buffer B (100nM NaCl and 500 mM imidazole in 50 mM Tris at pH 8.0). Purity of the sample was assessed using SDS PAGE analysis on Mini-PROTEAN Precast Gels (Bio-Rad). Fractions containing the protein of interest were then concentrated using Amicon Ultra centrifugal filters (10 kDa) and rinsed with a buffer containing Tris 50 mM, NaCl 100 mM, pH 8.

#### Protein-loaded P22 virus-like particles

The pBAD acceptor vector containing the cargo-SP fusion was co-transformed with pRSF-P22CP into *E. coli*, either Nico21(λDE3) or into BL21(λDE3) cells. Bacteria were grown on Luria-Bertani (LB) medium containing 100 μg/mL ampicillin and 50 μg/mL kanamycin at 37°C, 180 rpm until OD_600_ reached 0.6. Induction of the scaffold-fused protein was triggered by addition of L-arabinose at either 0.004% (low induction) or 0.2% L-arabinose (High induction) and temperature was lowered to 28°C. After 4 hours, induction of the coat protein vector was triggered by addition of isopropyl-β–D-1-thiogalactopyranoside (IPTG) to a final concentration of 1 mM. Cells were left over night and were harvested the next day by centrifugation at 5,000 g for 15 minutes, resuspended in phosphate buffered saline (PBS, pH 7.4) and lysed by passage through a homogenizer Emulsiflex-C5 (AVESTIN, Canada) five times at 20,000 PSI. The lysis was followed by centrifugation at 18,000 x g for 30 minutes to pellet the cellular debris. The supernatant was then loaded onto a gradient made of 3mL fractions of 50%, 40%, 30% and 20% Iodixanol in PBS, and subjected to ultracentrifugation for 2.5 hours at 150,000 g using either an SW 32 Ti or a SW41 Ti rotor. A band containing the P22 VLPs was extracted using a 1 mL syringe by side puncture, followed by desalting with a PD-Midi trap G-25 column (GE Healthcare Life Sciences). The constructs containing enzymes were desalted into 50 mM Tris, pH 8.0 with 50 mM NaCl as phosphate buffers have been found to inhibit some mevalonate kinases. For VLP preparations assayed for MK activity, a size exclusion purification step was performed using a S500 column HiPrep Sephacryl 16/60 S-500 HR (GE Healthcare Life Sciences), equilibrated with PBS buffer. Concentration of the VLPs was estimated by Bradford assay and densitometry using SDS PAGE analysis of serially diluted VLP preparations on Mini-PROTEAN Precast Gels (Bio-Rad), stained with GelCode Blue Stain (Thermo Fisher) and the average protein loading inside the VLP was estimated by densitometry using ImageJ [63] for at least three biological replicates.

#### Electron microscopy

Purified VLPs at ∼0.1 mg/mL were applied to formvar/carbon coated grids (ProSciTech Pty Ltd, Australia) and incubated for 2 minutes. Grids were then washed with 40 μL of distilled water for 30 seconds twice before staining with 2% uranyl acetate for 1 minute followed by blotted on filter paper. Images were taken either on a JEOL-1011 or a HITACHI HT7700 transmission electron microscope at accelerating voltage of 80 keV.

#### Enzyme assays

Mevalonate kinase activity was determined using a coupled assay as previously described [64]. (±)-Mevalonolactone was purchased from Sigma-Aldrich and dissolved in 50 mM Tris, pH 8.0 with 100 mM NaCl. Mevalonate was obtained by incubating mevalonolactone with KOH at a 1:1 molar ratio for 2 hours at 37°C, before re-adjusting the pH to 8.0. Estimation of the enzyme concentration in MK-VLPs was performed using Bradford (total protein) and densitometry (cargo component) and were performed with the various MK-VLPs and their free counterparts in 50 mM Tris, pH 8.0 with 50 mM NaCl. Reaction progress was monitored by following a coupled assay involving measuring NADH consumption at 340 nm on a UV spectrophotometer FLUOstar Omega (BMG Labtech) at 30°C. The coupled assay, adapted from previous publications [48,64], was performed using: 0-5 mM mevalonate, 5 mM ATP, 0.4 mM phosphoenol pyruvate, 5 units of pyruvate kinase, 5 units of lactate dehydrogenase and 0.2 mM NADH. Enzyme concentrations were between 11 nM and 64 nM depending on the MK and MK-VLP. Reactions were triggered by addition of a mevalonate and a non-limiting amount of 5 mM ATP was used. *K*_M_ and *k*_cat_ were calculated using Graphpad Prism 8 (www.graphpad.com). The Michaelis-Menten model was fitted to the data for the MKmaz and MKbur and the substrate inhibition model was fitted to the data in the case of MKvar. The Michaelis-Menten curves can be found in Supp Figures 2-4.

### Statistical analysis

Statistics and boxplots were prepared using the R software [65] and the package ggplot2 [66,67]. For each boxplot in figure 5, the bottom of the whiskers represent the minimum (calculated as Q1-1.5*interquartile range (IQR)) and the maximum (calculated as Q3+1.5*IQR), the bottom middle and top of the boxes represent the first quartile (Q1), median, third quartile (Q3) respectively, outliers are represented as dots. Analysis of Variance (ANOVA) was used to compare the different encapsulation parameters and two-tailed, unpaired Student’s T-test were used to compare the mean value of different VLP groups.

## ACKNOWLEDGEMENTS

This research was supported by a Research Grant from the Human Frontier Science Program (Ref.-No: RGP0012/2018). The work was performed on the traditional lands of the Yugarabul, Yuggera, Jagera and Turrbal peoples. Frank Sainsbury acknowledges support from the Commonwealth Scientific and Industrial Research Organisation (CSIRO) in the form of a Synthetic Biology Future Science Platform Fellowship. Donna McNeale was supported by a CSIRO Synbio PhD Top-up Scholarship. The authors acknowledge the facilities, and the scientific and technical assistance, of the Microscopy Australia Facility at the Centre for Microscopy and Microanalysis (CMM), The University of Queensland.

