## Supplementary Information for "Rapid design and prototyping of biocatalytic virus-like particle nanoreactors"

Contents:

**Supplementary Figure 1.** Loading density of MK cargo proteins in P22 VLPs.

**Supplementary Table 1.** ANOVA table analysing the impact of the various parameters on loading VLPs.

**Supplementary Figure 2.** Multiple sequence alignment of the three mevalonate kinases enzymes.

**Supplementary Figure 3.** Michaelis-menten curves for MKmaz and MKmaz-VLPs.

**Supplementary Figure 4.** Michaelis-menten curves for MKbur and MKbur-VLPs.

**Supplementary Figure 5.** Michaelis-menten curves for MKvar and MKvar-VLPs.

**Supplementary Table 2.** Kinetic parameters of the mevalonate kinase MKmaz.

**Supplementary Table 3.** Kinetic parameters of the mevalonate kinase MKbur.

**Supplementary Table 4.** Kinetic parameters of the mevalonate kinase MKvar.

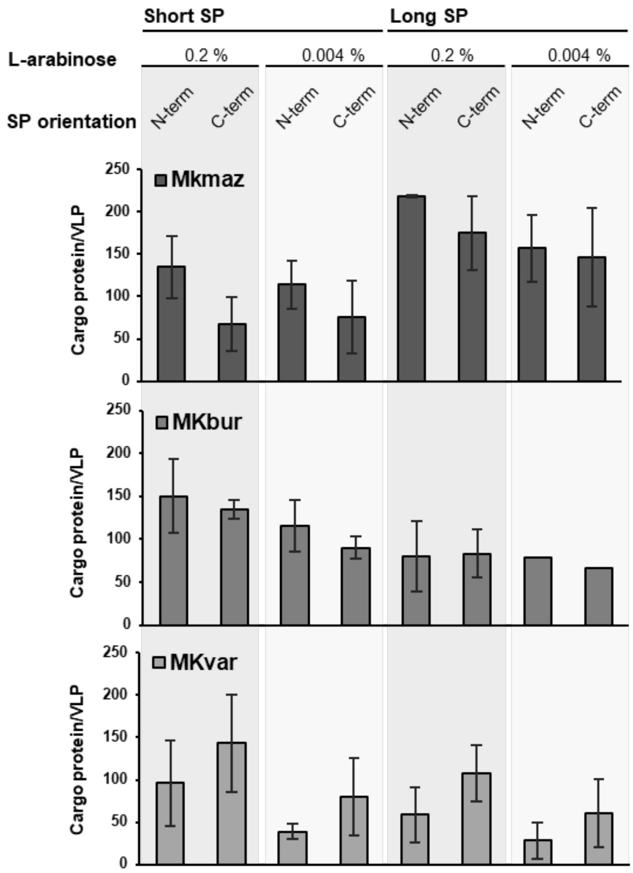


**Supplementary Figure 1. Loading density of MK cargo proteins in P22 VLPs.** SP = scaffold protein, C/N-term = fusion of the SP at the N-terminus or C-terminus of the cargo, percentage of L-arabinose corresponds to the concentration used to induce SP-cargo fusion gene expression. Densitometry of SDS-PAGE gels was used to estimate cargo loading (n=3).

**Supplementary Table 1. ANOVA table analysing the impact of the various parameters on loading VLPs.** (cargo protein identity, high/low arabinose, length of the scaffold, and orientation of the scaffold) loading ~ Cargo.protein*High.Low*Long.Short*N.C; where '***'is 0.001 '**' is 0.01 '*' is 0.05.

| | Sum of squares | Degree of freedom | F value | Pr (>F) P-value | |
|---|---|---|---|---|---|
| Cargo.protein | 29223 | 3 | 7.7559 | 0.0003519 | *** |
| High.Low | 29024 | 1 | 23.1086 | 0.000022973 | *** |
| Long.Short | 3 | 1 | 0.0025 | 0.9601262 | |
| N.C | 4192 | 1 | 3.3377 | 0.0753693 | |
| Cargo.protein:High.Low | 9499 | 3 | 2.5210 | 0.0719293 | . |
| Cargo.protein:Long.Short | 40585 | 2 | 16.1568 | 0.000007743 | *** |
| High.Low:Long.Short | 18 | 1 | 0.0142 | 0.9056277 | |
| Cargo.protein:N.C | 36208 | 3 | 9.6095 | 0.000070275 | *** |
| High.Low:N.C | 1664 | 1 | 1.3251 | 0.2566918 | |
| Long.Short:N.C | 435 | 1 | 0.3466 | 0.5594542 | |
| Cargo.protein:High.Low:Long.Short | 3394 | 2 | 1.3512 | 0.2707784 | |
| Cargo.protein:High.Low:N.C | 8295 | 3 | 2.2015 | 0.1032803 | |
| Cargo.protein:Long.Short:N.C | 457 | 2 | 0.1818 | 0.8344848 | |
| High.Low:Long.Short:N.C | 35 | 1 | 0.0282 | 0.8675293 | |
| Cargo.protein:High.Low:Long.Short:N.C | 18 | 2 | 0.0070 | 0.9930349 | |
| Residuals | 48983 | 39 | | | |

```
MKbur           M-----ITCSAPGKVYLFGEHAVVYGEPAICCAVDIRTRVTVSP------
MKmaz           M-----VSCSAPGKIYLFGEHAVVYGETAIACAVELRTRVRAEL------
MKvar           MSNLRHLRVSAPGKIILHGEHAVVYQKTAVALSLGLRTRLDLTETTDGRI
                *     :  *****: *.******* :.*:. :: :***:

MKbur           ---------------------ADTITISSSLGTTGIDFEV----------
MKmaz           ---------------------NDSITIQSQIGRTGLDFEK----------
MKvar           SIIMDKFLQHTSWSVEELSKIIDKVKIDANNPETELDQELVEDLRMMTSG
                                      *.:.*.:.   * :* *

MKbur           --------------------------HPYVSAVLERFQDISSFDGVDLRI
MKmaz           --------------------------HPYVSAVIEKMRKSIPINGVFLTV
MKvar           HHYQNGDGPAVGNPQAYSTQSVALVGFLYILVKLCKFSGKQRPPSIQISI
                                          . *: . : ::       .: : :

MKbur           SSDIPVGSGLGSSAAVTVATIKAMDTLLDLG---------------LELD
MKmaz           DSDIPVGSGLGSSAAVTIASIGALNELFGFG---------------LSLQ
MKvar           SSDIAISAGLGSSAAFAVCLSASLLSYLGIIVCDRKNCADVDGKLVPSAD
                .***.:.:*******.::.   ::   :.:                 . :

MKbur           D---IAKMGHEVEQNIQGTASPTDTYVCTMGGVVLIPQRKKLEL----ID
MKmaz           E---IAKLGHEIEIKVQGAASPTDTYVSTFGGVVTIPERRKLKT----PD
MKvar           QLALINHWAFMVEKIVHGSASGVDNAVSTYGGSIKYRN-NELTRIGSGLK
                :   * : .. :*  ::*:** .*. *.* ** :   : .:*       .

MKbur           CGILIGNTNIFSSTKELVGNVADLNERFPDVVGPVLSSIGKLSVI-----
MKmaz           CGIVIGDTGVFSSTKELVANVRQLRESYPDLIEPLMTSIGKISRI-----
MKvar           LDVLIVDTHVQRDTKKMLDIVRHRRKLYPAITNPVLEAIDGISETSSKIL
                 .::* :* :  .**:::  * . .: :* :  *:: :*. :*

MKbur           --GEGLVNDRDYVSVGELMNIDQGLLDAIGVSCAELSSLIYAARESGAYG
MKmaz           --GEQLVLSGDYASIGRLMNVNQGLLDALGVNILELSQLIYSARAAGAFG
MKvar           QHGDGLPTGEEYEVIADLVRMNQNLLSTLGVSHPKLDVICETASRFGQAG
                  *: *  . :*  :. *:.::*.**.::**.  :*. :  :*   *  *

MKbur           SKITGAGGGGCMVAISPREN--VDSVAE-------------AIGMAGGKV
MKmaz           AKITGAGGGGCMVALTAPEK--CNQVAE-------------AVAGAGGKV
MKvar           -KLTGAGGGGCAIVVLDPDMRQFEHLRESIIAEYRRMEFKPHLAELGGPG
                 *:******** :.:   :    : : *              :.  **

MKbur           VVANATDIGVRVECQ
MKmaz           TITKPTEQGLKVD--
MKvar           VLFHPVP--------
                .: :..
```

**Supplementary Figure 2. Multiple sequence alignment of the three mevalonate kinases enzymes**. Performed with CLUSTAL W (1.83), the orange section shows sections of MKvar not present in the two others and likely to lead to major structural differences.

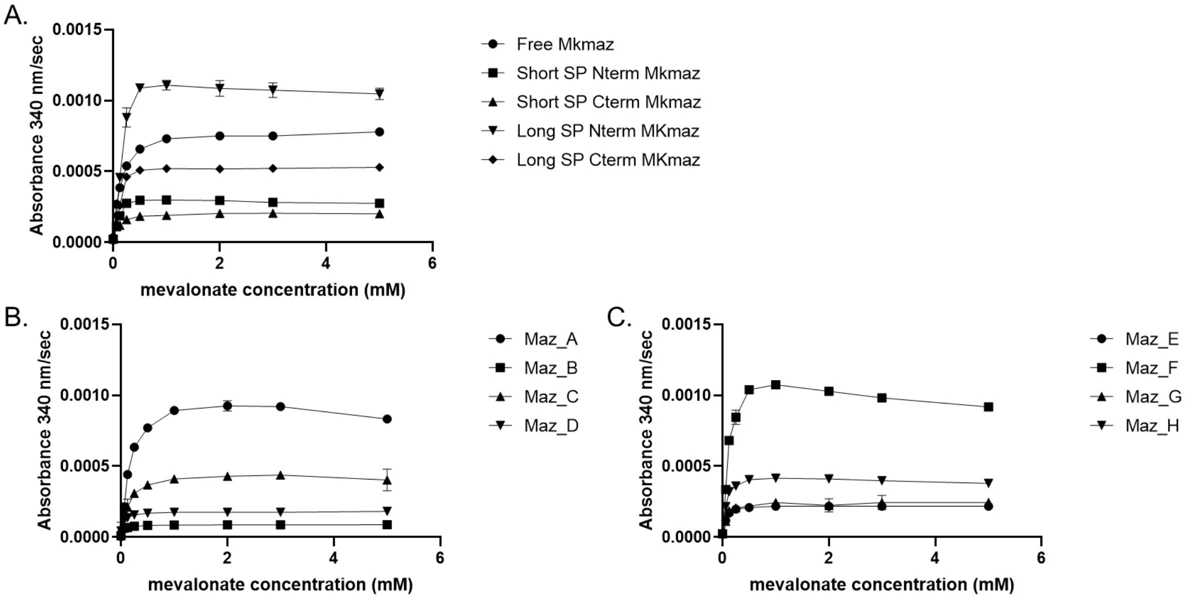


**Supplementary Figure 3. Michaelis-menten curves for MKmaz and MKmaz-VLPs.** A. free WT and fusions of MKmaz; B. MKmaz variants A-D loaded with the short SP N high arabinose, short SP C high arabinose short SP N low arabinose and short SP C low arabinose respectively; C. MKmaz variants E-H loaded with the long SP N high arabinose, long SP C high arabinose, long SP N low arabinose and long SP C low arabinose respectively.

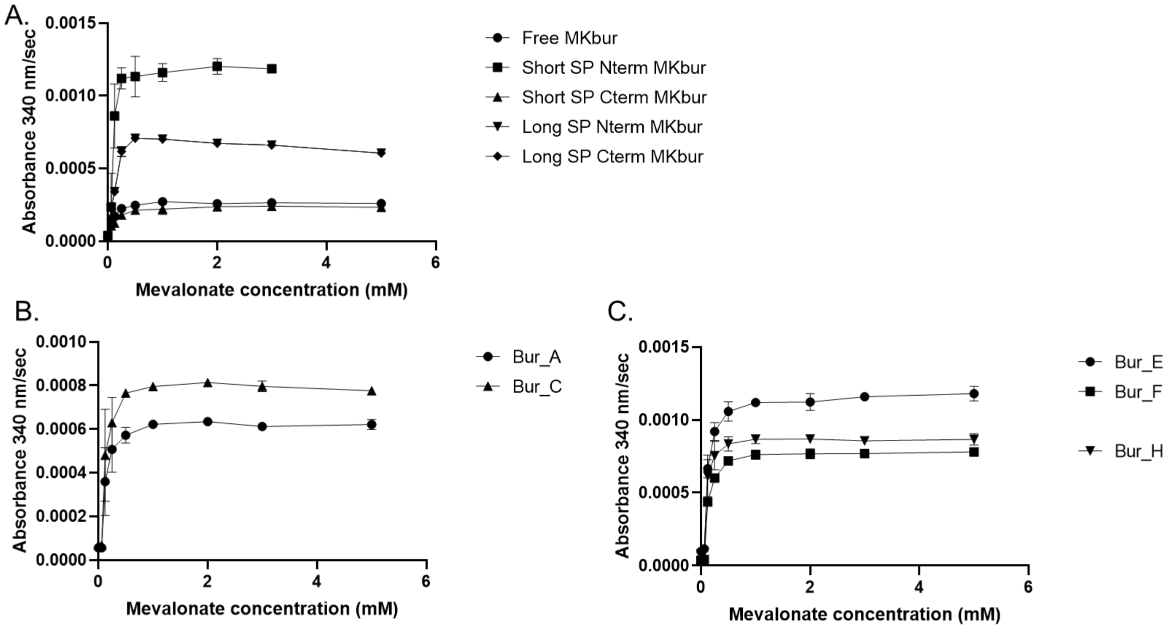


**Supplementary Figure 4. Michaelis-menten curves for MKbur and MKbur-VLPs.** A. free WT and fusions of MKbur; B. MKbur variants A-D loaded with the short SP N high arabinose, short SP C high arabinose short SP N low arabinose and short SP C low arabinose respectively; C. MKbur variants E-H loaded with the long SP N high arabinose, long SP C high arabinose, long SP N low arabinose and long SP C low arabinose respectively.

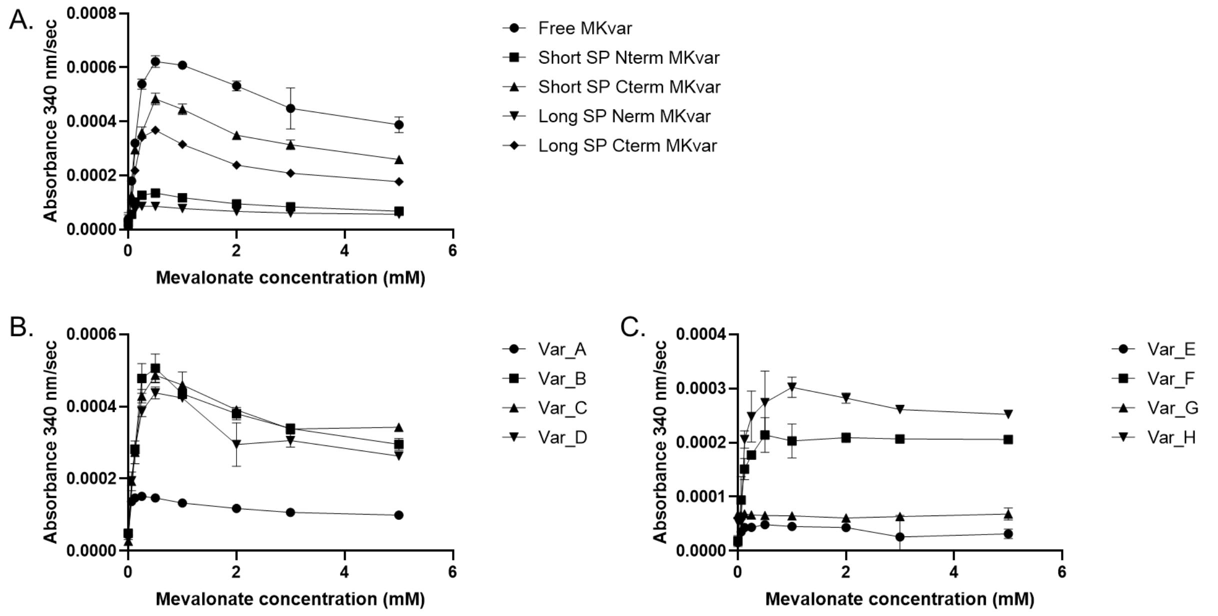


**Supplementary Figure 5. Michaelis-menten curves for MKvar and MKvar-VLPs.** A. free WT and fusions of MKvar; B. MKvar variants A-D loaded with the short SP N high arabinose, short SP C high arabinose short SP N low arabinose and short SP C low arabinose respectively; C. MKvar variants E-H loaded with the long SP N high arabinose, long SP C high arabinose, long SP N low arabinose and long SP C low arabinose respectively.

**Supplementary Table 2. Kinetic parameters of the mevalonate kinase MKmaz.** Apparent kinetic parameters of free enzyme and seven nanoreactors encapsulating MKmaz at different loading densities (data represent averages of n=4 technical replicates ± stdev). High and Low refer to different expression induction levels of cargo enzymes for VLP loading. Bold lines indicate low-loading examples

| | | | app $K_M$ (mM) | app $k_{cat}$ ($s^{-1}$) | app $k_{cat}$ / app $K_M$ | Loading |
|---|---|---|---|---|---|---|
| | | untagged | 0.12 ±0.005 | 10.1 ±0.08 | 82 ±17 | - |
| Short SP | | N- | 0.07 ±0.009 | 3.6 ±0.08 | 50 ±9 | - |
| | | C- | 0.07 ±0.008 | 1.7 ±0.03 | 24 ±4 | - |
| Long SP | | N- | 0.15 ±0.019 | 7.3 ±0.022 | 50 ±11 | - |
| | | C- | 0.12 ±0.013 | 3.5 ±0.08 | 30 ±6 | - |
| Short SP | High | N- | 0.14 ±0.012 | 4.6 ±0.08 | 32 ±7 | 175 |
| | High | C- | 0.05 ±0.003 | 1.0 ±0.01 | 22 ±2 | 48 |
| | Low | N- | 0.11 ±0.017 | 2.1 ±0.06 | 20 ±4 | 56 |
| | Low | C- | 0.03 ±0.007 | 0.9 ±0.02 | 26 ±3 | 14 |
| Long SP | High | N- | 0.05 ±0.003 | 1.6 ±0.02 | 32 ±3 | 218 |
| | High | C- | 0.08 ±0.01 | 2.9 ±0.02 | 21 ±9 | 206 |
| | Low | N- | 0.06 ±0.02 | 1.9 ±0.1 | 20 ±20 | 185 |
| Long SP | Low | C- | 0.05 ±0.004 | 1.9 ±0.03 | 26 ±5 | 188 |
| | **High** | **N-** | **0.15 ±0.015** | **7.2 ±0.15** | **49±8** | **55** |
| | **High** | **C-** | **0.07 ±0.008** | **2.4 ±0.05** | **32±7** | **28** |

**Supplementary Table 3. Kinetic parameters of the mevalonate kinase MKbur.** Apparent kinetic parameters of free enzyme and five VLP nanoreactors encapsulating MKbur at different loading densities (data represent averages of n=4 technical replicates ± stdev). High and Low refer to different expression induction levels of cargo enzymes for VLP loading.

| | | | app $K_M$ (mM) | app $k_{cat}$ ($s^{-1}$) | app $k_{cat}$ / app $K_M$ | loading |
|---|---|---|---|---|---|---|
| | | untagged | 0.06 ±0.005 | 4.8 ±0.08 | 88 ± 13 | - |
| Short SP | | N- | 0.09 ±0.019 | 6.3 ±0.28 | 66 ±14 | - |
| | | C- | 0.09 ±0.012 | 4.6 ±0.11 | 48 ±9 | - |
| Long SP | | N- | 0.15 ±0.019 | 6.6 ±0.19 | 45 ±10 | - |
| | | C- | 0.09 ±0.014 | 3.2 ±0.09 | 35 ±7 | - |
| Short SP | High | N- | 0.14 ±0.025 | 3.3 ±0.13 | 24 ±5 | 105 |
| | Low | N- | 0.14 ±0.025 | 4.2 ±0.17 | 31 ±7 | 85 |
| Long SP | High | N- | 0.14 ±0.019 | 6.1 ±0.18 | 43 ±9 | 34 |
| | High | C- | 0.15 ±0.021 | 4.1 ±0.13 | 27 ±6 | 53 |
| | Low | N- | 0.11 ±0.020 | 4.5 ±0.17 | 42±9 | 67 |

**Supplementary Table 4. Kinetic parameters of the mevalonate kinase MKvar.** Apparent kinetic parametesr of free enzyme and seven VLP nanoreactors encapsulating MKvar at different loading densities (data represent averages of n=4 technical replicates ± stdev). High and Low refer to different expression induction levels of cargo enzymes for VLP loading.

| | | | app $K_M$ (mM) | app $k_{cat}$ ($s^{-1}$) | $K_i$ (mM) | app $k_{cat}$ / app $K_M$ | Loading |
|---|---|---|---|---|---|---|---|
| | | untagged | 0.24 ±0.04 | 21.9 ±0.40 | 2.7 ±0.4 | 92 ±11 | - |
| Short SP | | N- | 0.12 ±0.03 | 1.9 ±0.02 | 0.9 ±0.5 | 8 ±1 | - |
| | | C- | 0.23 ±0.04 | 3.7 ±0.07 | 3.7 ±0.4 | 16 ±2 | - |
| Long SP | | N- | 0.05 ±0.01 | 0.9 ±0.01 | 3.9 ±1 | 16 ±1 | - |
| | | C- | 0.25 ±0.05 | 1.9 ±0.05 | 1 ±0.3 | 8 ±1 | - |
| Short SP | High | N- | 0.09 ±0.02 | 5.1 ±0.09 | 5.1 ±1.2 | 58 ±5 | 86 |
| | High | C- | 0.14 ±0.03 | 5.6 ±0.14 | 3.0 ±0.7 | 42 ±4 | 117 |
| | Low | N- | 0.14 ±0.02 | 3.1 ±0.05 | 4.2 ±0.8 | 22 ±2 | 50 |
| | Low | C- | 0.12 ±0.02 | 2.9 ±0.06 | 3.1 ±0.6 | 23 ±2 | 44 |
| Long SP | High | N- | 0.03 ±0.03 | 0.2 ±0.03 | 5.1 ±9.7 | 6 ±1 | 39 |
| | High | C- | 0.08 ±0.02 | 0.5 ±0.02 | 31.8 ±3.1 | 6 ±1 | 91 |
| | Low | C- | 0.16 ±0.04 | 0.8 ±0.04 | 8.4 ±9 | 5 ±1 | 57 |
