## Supplementary Data for "Rapid design and prototyping of biocatalytic virus-like particle nanoreactors"

**Supplementary Data.** Description and sequences of the plasmids created for P22 cargo loading. Double click .gb file icon for full annotated sequence.

| **Name** | **Function** | **Origin** | **Induction** | **Selection** | **Sequence (.gb)** |
| --- | --- | --- | --- | --- | --- |
| pBAD-HisN | Acceptor vector for cargo expressed with N-terminal 6xHis-tag only | ColEI | Arabinose | Ampicillin | 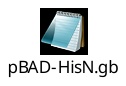 |
| pBAD-HisC | Acceptor vector for cargo expressed with C-terminal 6xHis-tag only | ColEI | Arabinose | Ampicillin | 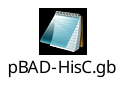 |
| pBAD-P22-shSPN | Acceptor vector for cargo loading with N-terminal short SP^a^ | ColEI | Arabinose | Ampicillin | 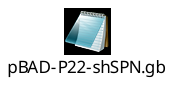 |
| pBAD-P22-shSPC | Acceptor vector for cargo loading with C-terminal short SP^a^ | ColEI | Arabinose | Ampicillin | 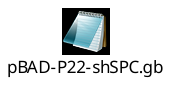 |
| pBAD-P22-lgSPN | Acceptor vector for cargo loading with N-terminal long SP^a^ | ColEI | Arabinose | Ampicillin | 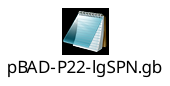 |
| pBAD-P22-lgSPC | Acceptor vector for cargo loading with C-terminal long SP^a^ | ColEI | Arabinose | Ampicillin | 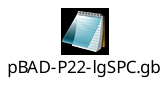 |
| pRSF-P22CP | P22 coat protein (CP) expression vector | RSF | IPTG | Kanamycin | 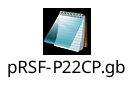 |
| pRSF-P22-shSPN-CP | Acceptor vector for cargo loading with N-terminal short SP | RSF | IPTG^b^ | Kanamycin | 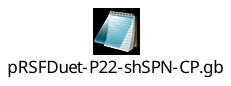 |
| pRSF-P22-shSPC-CP | Acceptor vector for cargo loading with C-terminal long SP | RSF | IPTG^b^ | Kanamycin | 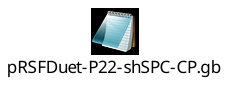 |
| pTwist-YFP-donor | Example donor vector/ordering template | ColEI | N/A | Chloramphenicol | 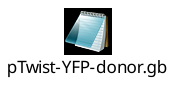 |

a) **co-expression with a pRSF vector expressing CP is required for encapsulation**
b) **IPTG simultaneously induces both the CP and SPfusion cargo**
